# Uncovering the roles of the scaffolding protein CsoS2 in mediating the assembly and shape of the α-carboxysome shell

**DOI:** 10.1101/2024.05.14.594188

**Authors:** Tianpei Li, Taiyu Chen, Ping Chang, Xingwu Ge, Vincent Chriscoli, Gregory F. Dykes, Qiang Wang, Lu-Ning Liu

## Abstract

Carboxysomes are proteinaceous organelles featuring icosahedral protein shells that enclose the carbon-fixing enzymes, Rubisco, alone with carbonic anhydrase. The intrinsically disordered scaffolding protein CsoS2 plays a vital role in the construction of α-carboxysomes through bridging the shell and cargo enzymes. The N-terminal domain of CsoS2 binds Rubisco and facilitates Rubisco packaging within the α-carboxysome, whereas the C-terminal domain of CsoS2 (CsoS2-C) anchors to the shell and promotes shell assembly. However, the role of the middle region of CsoS2 (CsoS2-M) has remained elusive. Here, we conducted indepth examinations on the function of CsoS2-M in the assembly of the α-carboxysome shell by generating a series of recombinant shell variants in the absence of cargos. Our results reveal that CsoS2-M assists CsoS2-C in the assembly of the α-carboxysome shell and plays an important role in shaping the α-carboxysome shell through enhancing the association of shell proteins on both the facet-facet interfaces and flat shell facets. Moreover, CsoS2-M is responsible for recruiting the C-terminal truncated isoform of CsoS2, CsoS2A, into α-carboxysomes, which is crucial for Rubisco encapsulation and packaging. This study not only deepens our knowledge of how the carboxysome shell is constructed and regulated but also lays the groundwork for engineering and repurposing carboxysome-based nanostructures for diverse biotechnological purposes.

## Introduction

Carboxysomes (CBs) are specialized organelle-like proteinaceous microcompartments ubiquitous in cyanobacteria and some proteobacteria, playing a pivotal role in CO_2_ fixation (Kerfeld and Melnicki, 2016; Liu, 2022). Diverging from eukaryotic counterparts, CBs are entirely proteinaceous, featuring a polyhedral shell and cargo enzymes crucial for CO_2_ fixation (Yeates et al., 2008). The proteinaceous shell comprises primarily three groups of building blocks, including hexamers and trimers that form the facets (Tsai et al., 2007; Klein et al., 2009; Larsson et al., 2017), and pentamers that cap the vertices (Cai et al., 2009; Zhao et al., 2019). In addition, the scaffolding proteins bridge the shell and cargo enzymes, mediating the assembly of CBs (Cai et al., 2015; Wang et al., 2019; Oltrogge et al., 2020; Zang et al., 2021). Through self-assembly *in vivo*, thousands of these building blocks form a highly ordered icosahedral shell, sequestering ribulose-1,5-bisphosphate carboxylase oxygenase (Rubisco) and carbonic anhydrase for the construction of a functional entity (Liu, 2022). Moreover, the porous shell functions as a barrier, selectively modulating the influx and efflux of metabolites, significantly facilitating the catalytic performance of the interior enzymes (Dou et al., 2008; Klein et al., 2009; Menon et al., 2010; Mahinthichaichan et al., 2018; Faulkner et al., 2020; Sarkar D, 2024). The self-assembly, permeability, and catalytic enhancement properties of CBs make them an appealing bioengineering target for applications in crop engineering, biofuel production, metabolic enhancement, and therapeutics (Li et al., 2020; Borden and Savage, 2021; Chen et al., 2023; Jiang et al., 2023).

Based on the Rubisco phylogeny, CBs are classified into two categories: α-CBs that are primarily encoded by the *cso* operon and β-CBs that are primarily encoded by the *ccm* operon (Rae et al., 2013; Turmo et al., 2017). Unlike β-CBs that undertake a “Cargo first” assembly pathway (Cameron et al., 2013; Chen et al., 2013), the self-assembly of α-CBs is presumed to follow a “Shell first” or “Concomitant shell−core assembly” mode (Menon et al., 2008; Iancu et al., 2010; Dai et al., 2018). This unique assembly pathway offers α-CBs with greater potential to engineer empty shell structures that can be used to enclose foreign cargos and molecules for generation of new nanobioreactors and scaffolding nanomaterials. Previous studies have demonstrated the possibilities of engineering intact α-CBs (Bonacci et al., 2012; Flamholz et al., 2020; Sun et al., 2022), entire or simplified α-CB shells (Li et al., 2020; Tan et al., 2021; Huang et al., 2022; Ni et al., 2023; Li et al., 2024), and α-CB-based nanobioreactors in *Escherichia coli* (*E. coli*) (Li et al., 2020; Jiang et al., 2023), as well as transforming α-CBs into plant chloroplasts for boosting carbon fixation (Long et al., 2018; Chen et al., 2023). In this context, understanding the exact structural organization and assembly mechanisms of α-CBs and α-CB shells is fundamental for rational design and reprogramming of CB-inspired nanostructures.

Recent studies using protein crystallization and cryo-electron microscopy (cryo-EM) have provided valuable insights into the building principles of α-CBs, in particular the crucial role of the scaffolding protein CsoS2 in orchestrating the assembly of shell proteins and cargos (Oltrogge et al., 2020; Tan et al., 2021; Metskas et al., 2022; Ni et al., 2022; Evans et al., 2023; Zhou et al., 2024). CsoS2 comprises three distinct regions: an N-terminal region (CsoS2-N), a middle region (CsoS2-M), and a C-terminal region (CsoS2-C) (Figure 1a). Moreover, each region contains various repetitive fragments: four N-repeats in CsoS2-N, six M-repeats in CsoS2-M, and three C-repeats in CsoS2-C. It has been demonstrated that CsoS2-N binds with Rubisco through multivalent interactions, playing a role in recruiting Rubisco (Oltrogge et al., 2020; Ni et al., 2022); whereas CsoS2-C binds with shell proteins via its C-repeats, functioning as a “molecular thread” to stitch multiple shell proteins, thereby facilitating shell assembly (Ni et al., 2023). CsoS2-M has been suggested to be important in determining the size of α-CBs (Oltrogge et al., 2024). In addition, a recent study reveals that CsoS2-M binds to multiple hexamers on the shell facets through multivalent interactions in α-CBs from the marine cyanobacterium *Prochlorococcus* MED4, implying an important role of CsoS2-M in shaping the structure of α-CBs (Zhou et al., 2024).

**Figure 1.**
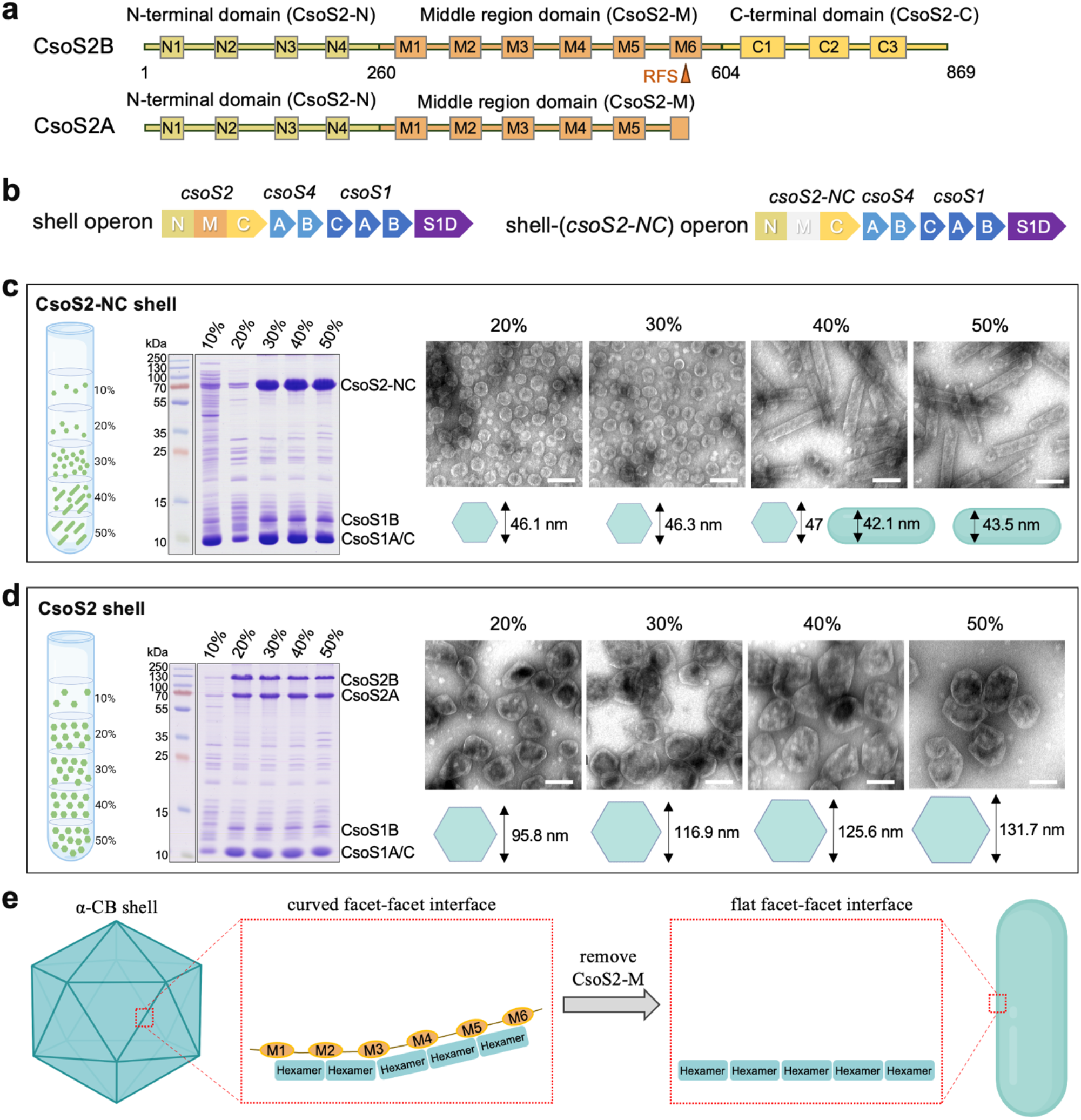
CsoS2-M defines the size and shape of the α-CB shells. **(a)** The domain arrangement of CsoS2. The N-terminal (CsoS2-N), Middle (CsoS2-M) and C-terminal (CsoS2-C) domains are colored sand, orange and yellow, respectively. A ribosomal frameshift site (RFS) located at the 6^th^ repeat (M6) of CsoS2-M leads to the production of a short CsoS2A isoform. **(b)** The genetic arrangements of shell and shell-(*csoS2-NC*) operons, with the deleted region colored grey. **(c-d)** Cartoon models showing the distribution of CsoS2-NC shell and tubular assemblies (c), and CsoS2 shells (with unmodified CsoS2) (d) across 10−50% sucrose fractions after ultracentrifugation (left panel). SDS-PAGE of CsoS2-NC shell and tubular assemblies (c) and CsoS2 shells (d) isolated from 10−50% sucrose fractions (middle panel). Transmission EM images displaying CsoS2-NC shell and tubular assemblies (c) and CsoS2 shells (d) in 20-50% sucrose fractions, respectively (right panel). The average diameters of shells and widths of tubular structures in various sucrose fractions are depicted below the EM images. Scale bar: 100 nm. **(e)** A cartoon model illustrating a possible mechanism by which CsoS2-M defines the shell curvature. CsoS2-M is likely involved in adjusting the tilt angles between neighbouring shell facets by enhancing the association of shell proteins on the facet-facet interfaces.

CsoS2 in many microorganisms, including the model chemoautotrophic bacterium *Halothiobacillus neapolitanus* (*H. neapolitanus*), contains a ribosomal frameshift site (RFS) within CsoS2-M, leading to the premature termination and production of a full-length isoform CsoS2B, and a shorter isoform CsoS2A (Chaijarasphong et al., 2016) (Figure 1a). Compared to CsoS2B, CsoS2A lacks CsoS2-C that is crucial for binding with the α-CB shell. How CsoS2A is incorporated within the α-CB, how it coordinates with CsoS2B, as well as the precise roles of CsoS2-M and CsoS2A in determining shell formation and architecture remains elusive.

Here, by generating a series of recombinant α-CBs shells derived from *H. neapolitanus*, we systematically evaluated the roles of individual domains of CsoS2, without cargo proteins, in modulating the shell formation. We show that CsoS2-M assists CsoS2-C in enhancing the connections between hexamers that are distal from the shell vertices, thereby playing a dominant role in determining the size and shape of α-CB shells. These findings enable us to develop a model to elucidate the mechanisms by which CsoS2 facilitates the assembly of the α-CB shell. This study advances our knowledge of the self-assembly and structural basis of α-CBs, which lays the groundwork for future engineering and refinement of α-CBs for various biotechnological and biomedical applications.

## Results

### CsoS2-M plays a role in determining the shell size and morphology

In previous work, we have constructed recombinant α-CB shells with an average size of ∼120 nm in *E. coli*, by expressing a shell operon derived from *H. neapolitanus* (Li et al., 2020). This shell operon consists of genes encoding CsoS2, pentamers CsoS4A/4B, hexamers CsoS1A/1B/1C, and trimers CsoS1D (Figure 1b). To investigate the role of CsoS2-M during shell assembly, we generated a shell-(*csoS2*-*NC*) operon by deleting only the nucleotide sequences encoding CsoS2-M from the shell operon (Figure 1b). The shell-(*csoS2*-*NC*) operon was expressed in *E. coli* and the resulting shell assemblies were purified by sucrose gradient centrifugation. SDS-polyacrylamide gel electrophoresis (SDS-PAGE) confirmed the presence of CsoS2-NC and CsoS1A/B/C in various sucrose fractions, with a predominant distribution in the 30−50% sucrose fractions (Figure 1c). Surprisingly, despite the same protein composition in the 20−50% sucrose fractions identified by SDS-PAGE, EM revealed distinct tubular structures with a mean width of ∼42 or ∼44 nm in 40% and 50% sucrose fractions, whereas polyhedral shells (∼46 nm on average in diameter) were enriched in 20% and 30% sucrose fractions (Figure 1c). By contrast, expression of the shell operon in *E. coli* exclusively produced polyhedral shells, despite that the shell size varies among different sucrose fractions (Figure 1d). Moreover, the polyhedral shells (∼46 nm) produced by the shell-(*csoS2*-*NC*) operon were remarkably smaller than the intact α-CB shells (∼120 nm) (Figure 1c, 1d). These results indicate that CsoS2-M plays an important role in defining the size and shape of α-CB shells.

We speculate that CsoS2-M is involved in adjusting the tilt angles between neighbouring facets of the icosahedral shell (Figure 1e). Without CsoS2-M, the interaction between hexamers situated at the curved facet-facet interface is diminished, resulting in the formation of a flat facet-facet interface and eventually the generation of low-curvature tubular structures (Figure 1e). Consistently, the cryo-EM structure of the simple and small α-CB from *Prochlorococcus* revealed that CsoS2-M binds to multiple hexamers distributed on the adjacent facets of the icosahedral shell through multivalent interactions (Zhou et al., 2024).

### Roles of CsoS2-N and CsoS2-C in the shell assembly

We also investigated the role of CsoS2-N and CsoS2-C by deleting *csoS2-N* and *csoS2-C* from the shell-(*csoS2*-*NC*) operon (Figure 2a). Removing CsoS2-N alone (shell-(*csoS2-C*) operon) or both CsoS2-N and CsoS2-C (shell-(Δ*csoS2*) operon) resulted in the production of polyhedral shells exclusively (Figure 2b). The CsoS2-C shells were ∼41 nm in diameter, slightly smaller than the CsoS2-NC shells (∼46 nm), which implies that CsoS2-N has no substantial effect on the shell size. This supports the finding that CsoS2-N interacts with Rubisco to facilitate Rubisco encapsulation into the α-CB (Oltrogge et al., 2020). By contrast, the Δ*csoS2* shells (∼23 nm) were only half the size of the CsoS2-C shells (∼41 nm) (Figure 2b, Figure S1), suggesting that CsoS2-C has a significant impact on the shell size through interactions with shell proteins. In line with this, our recent work has shown that the binding of CsoS2-C with shell proteins could result in an increase in the size of “mini-shells”, which are made up of CsoS1A hexamers and CsoS4A pentamers, from 25 nm to 37 nm (Ni et al., 2023).

**Figure 2.**
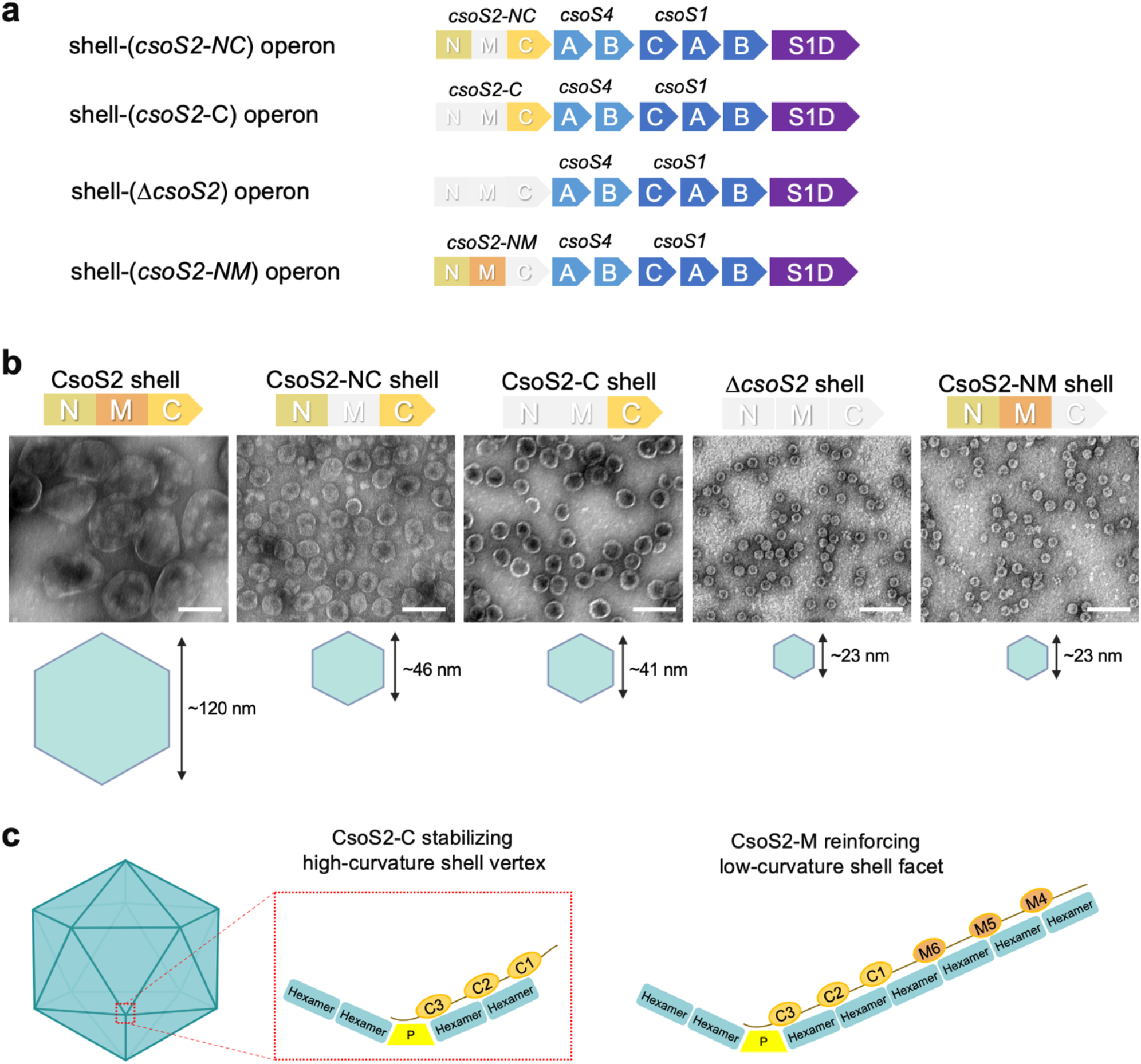
The role of individual domains of CsoS2 in modulating shell size. **(a)** The genetic arrangements of shell-(*csoS2-NC*), shell-(*csoS2-C*), shell-(Δ*csoS2*) and shell-(*csoS2-NM*) operons. **(b)**. Transmission EM images of CsoS2 shells, CsoS2-NC shells, CsoS2-C shells, Δ*csoS2* shells, and CsoS2-NM shells, respectively. The average sizes are displayed below the EM image. Scale bar: 100 nm. **(c)**. A cartoon model depicting interactions of CsoS2-M and CsoS2-C with shell proteins during shell assembly; CsoS2-M assists CsoS2-C in enhancing the connections between the hexamers that are distal from the shell vertices. P, pentamer.

To further evaluate the role of CsoS2-C in modulating the shell size, we also generated a shell-(*csoS2-NM*) operon by deleting *csoS2-C* from the shell operon (Figure 2a). Interestingly, expression of the shell-(*csoS2-NM*) operon in *E. coli* yielded exclusively mini-shells with diameters of ∼23 nm (Figure 2b). This result confirms the importance of CsoS2-C in forming shells with the diameter larger than ∼23 nm and suggests that CsoS2-M alone is insufficient to drive the assembly of intact α-CB shells (∼120 nm).

Given that CsoS2-C binds to the high-curvature pentamer-hexamer and hexamer-hexamer interfaces (Ni et al., 2023), it is likely that the multivalent interactions between CsoS2-C and shell proteins close to shell vertices are essential for the formation of α-CB shells and may potentially initiate the assembly of the α-CB shell; subsequently, CsoS2-M assists CsoS2-C by strengthening the low-curvature hexamer-hexamer interfaces that are distant from the shell vertices, resulting in the formation of intact α-CB shells (Figure 2c).

### The repeating motifs of CsoS2-M define the shell size

CsoS2-M contains six repeating fragments (M1–M6), each of which is ∼50 residues in length, separated by short linker sequences of 5∼15 residues, and possesses three conserved [V/I/M][T/S]G motifs (Figure 1a, Figure S2). To delineate in-depth how CsoS2-M regulates the shell size and shape, we generated a series of CsoS2 and CsoS2B variants with varying numbers of M-repeats in the M-region (Figure 3a, 3b). All the CsoS2 variants retained the RFS at the 6^th^ M-repeat (M6), allowing the production of both CsoS2B and CsoS2A isoforms, each with a varying number of M-repeats. By contrast, the CsoS2B variants lack M6, leading to the exclusive production of the CsoS2B isoform with a varying count of M-repeats. Given the high conservation among different M-repeats, individual differences in M-repeats were not considered in the design of these CsoS2 and CsoS2B variants (Figure S2).

**Figure 3.**
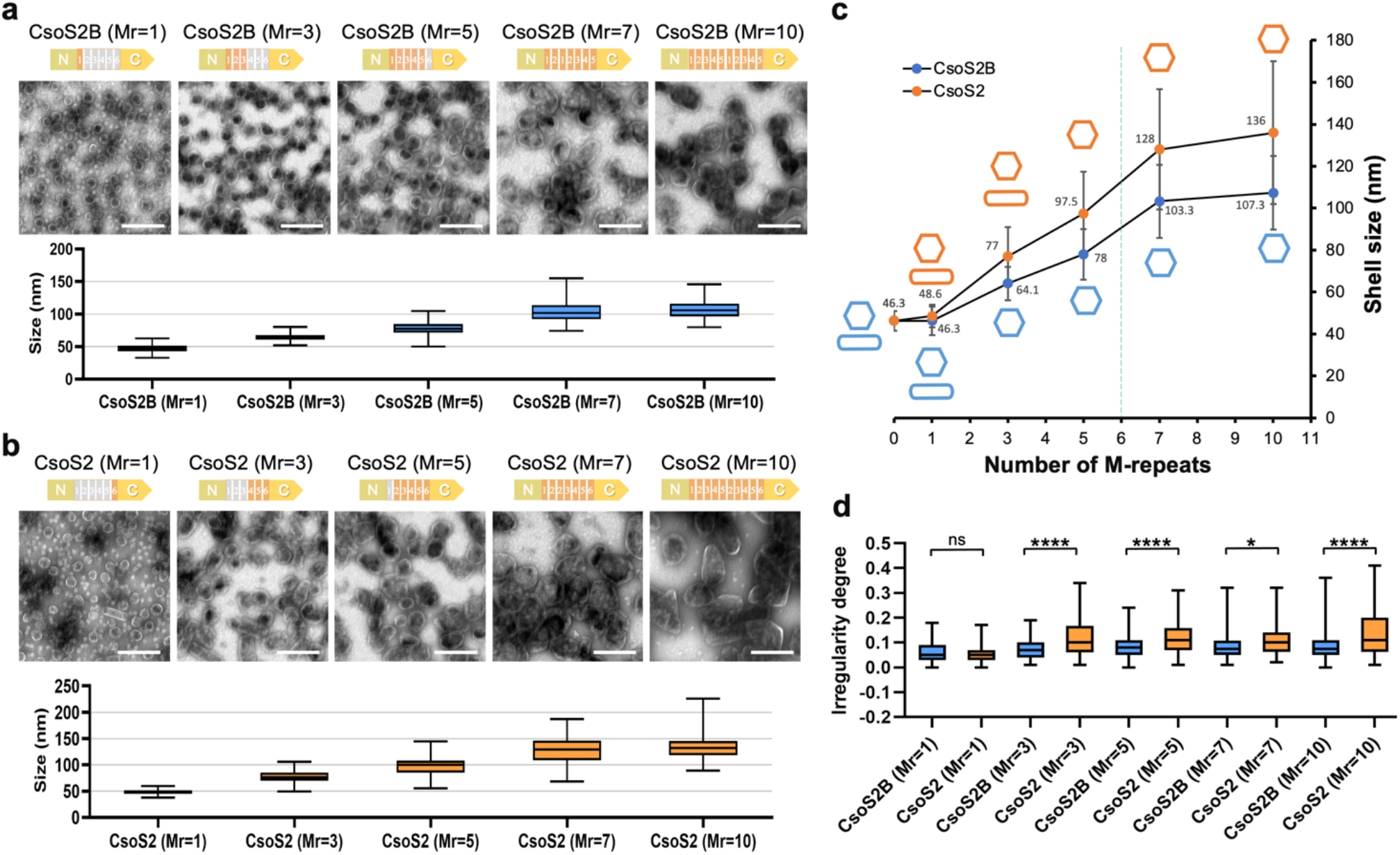
The number of M-repeats in CsoS2-M determines the shell size and shape. **(a)** A series of CsoS2B shell variants with M-repeats range from 1 (Mr=1) to 10 (Mr=10) in CsoS2B. Transmission EM images of each shell variant are displayed above the size distribution plot. Scale bar: 200 nm. Samples were collected from the sucrose fraction enriched with the most abundant shell proteins, as determined by SDS-PAGE (Figure S3). **(b)** A series of CsoS2 shell variants with M-repeats range from 1 (Mr=1) to 10 (Mr=10) in CsoS2. Scale bar: 200 nm. **(c)** Plot of the mean diameter of various shell variants as a function of the number of M-repeats. The cartoon models colored orange or blue represent the morphology of CsoS2 or CsoS2B shell variants, respectively. Native CsoS2 contains 6 M-repeats, as indicated by the green dashed line, which are of biologically importance for the formation of large polyhedral α-CB structures. **(d)** The irregularity degree of CsoS2B and CsoS2 shell variants. *, 0.01 ≤ *p* ≤ 0.05; ****, *p* ≤ 0.0001; ns, no significance (n = 100, two-tailed unpaired t-test). Box plots indicate the median (middle line in the box), 25^th^ percentile (bottom line of the box), and 75^th^ percentile (top line of the box).

The resulting shell variants were purified from *E. coli* using sucrose gradient centrifugation. SDS-PAGE verified the presence of CsoS2B variants in all the CsoS2B shell variants, and the coexistence of CsoS2B and CsoS2A isoforms in the CsoS2 shell variants, along with the main shell proteins CsoS1A/B/C (Figure S3). An exception is the CsoS2 shell variant that contains only a single <u>M</u>-repeat (M6), termed CsoS2 (<u>Mr</u>=1) shell, in which only CsoS2B(M6) was incorporated in the purified shell (Figure S3b). This suggests that the short isoform CsoS2A(M6) with a single M-repeat was unable to be encapsulated within the shell. EM further revealed that the size of the CsoS2B shell variants exhibited a significant increase, growing from ∼46 nm to ∼107 nm, corresponding to the rise in the number of M-repeats from one (Mr=1) to ten (Mr=10) (Figure 3a). Similarly, the size of the CsoS2 shell variants increased from ∼49 nm to ∼136 nm (Figure 3b). Interestingly, this upward trajectory seemed to decelerate when the number of M-repeats in both CsoS2 and CsoS2B shells exceeded seven (Mr=7) (Figure 3c). Overall, our results indicate that the number of CsoS2-M repeats is a key factor in determining the size of α-CB shells; larger shells are formed when the number of M-repeats increases. These findings support our hypothesis that CsoS2-M plays a role in reinforcing hexamer-hexamer interfaces that are distant from the shell vertices. More M-repeats accommodated in CsoS2-M will lead to the formation of larger shell facets and stable facet-facet interfaces, essential for the assembly of large polyhedral shells (Figure 2c, Figure S4). It is worth mentioning that the steady increase in the number of M-repeats did not lead to an infinite expansion of empty shells, suggesting that there should be other factors involved in defining the shell size. Consistent with our observations, an increase in the size of recombinant α-CBs that contain cargos was also found as the number of M-repeats increased (Oltrogge et al., 2024).

We also found that the CsoS2 shell variants (containing both CsoS2A and CsoS2B) appeared to be greater in size than the CsoS2B shells with the same number of M-repeats, suggesting that CsoS2A plays a role in facilitating the assembly of larger shells (Figure 3c). This is consistent with the previous finding that recombinant α-CBs, with cargos, containing both CsoS2 isoforms were larger than those with only CsoS2B (Oltrogge et al., 2024). Moreover, CsoS2 (Mr=3) featuring three M-repeats produced both polyhedral shells and tubular structures in the heavier sucrose fraction, whereas CsoS2B (Mr=3) with three M-repeats yielded exclusively polyhedral shells (Figure 3c, S4b). The resulting CsoS2 shells also displayed greater heterogeneity compared to the CsoS2B shells with the same number of M-repeats (Figure 3d). These results suggest that CsoS2A is involved in governing both the shell size and curvature. The additional M-repeats of CsoS2A, which are absent in the CsoS2B shells, might coordinate with the M-repeats of CsoS2B to facilitate the assembly of larger shells. This is further supported by the observations that the CsoS2 (Mr=1) shell and CsoS2B (Mr=1) shell, both of which have no CsoS2A, exhibit similar shell sizes (Figure 3c, Figure S3a, 3b). On the other hand, without CsoS2-C for anchoring near the shell vertices, CsoS2A might be able to adopt more flexible binding modes with shell proteins, which leads to the higher structural heterogeneity of the CsoS2A-containing shells.

### Encapsulation of CsoS2A is independent of its interaction with CsoS2-M of CsoS2B

It was assumed that the interactions between the CsoS2-M of CsoS2A and CsoS2B ensure the recruitment of CsoS2A into the α-CB (Cai et al., 2015). To test this, we constructed a plasmid to express CsoS2A with enhanced green fluorescent protein (GFP) fused at its N-terminus (GFP-CsoS2A), and fused mCherry to the C-terminus of CsoS1B (CsoS1B-mCherry) in those shell constructs. This allowed us to visualize the distribution of CsoS2A and shell variants in *E. coli*. Additionally, to eliminate the influence of endogenous CsoS2A, we constructed the shell-(*csoS2B-only*) operon by removing the RFS of *csoS2* (Chaijarasphong et al., 2016; Oltrogge et al., 2024), to express CsoS2B only without CsoS2A, along with shell proteins. The formation of the CsoS2B-only shells was confirmed by SDS-PAGE and EM (Figure S5). The CsoS2B-only shells labeled with mCherry were used for the following fluorescence co-localization analysis.

These mCherry-labeled shells, including the CsoS2 shells (with unmodified CsoS2), CsoS2B-only shells, CsoS2-NC shells, and CsoS2-C shells, were either expressed individually or co-expressed with GFP-CsoS2A. Confocal images of *E. coli* cells expressing different mCherry-labeled shell variants showed similar polar distribution of shell assemblies (Figure 4a). Intriguingly, colocalization of GFP-CsoS2A (green) and CsoS1B-mCherry (red) was visualized in cells expressing the CsoS2 shells, CsoS2B-only shells, CsoS2-NC shells, or CsoS2-C shells (Figure 4b). These data indicate that CsoS2A could be incorporated into the shells without the assistance of endogenous CsoS2-M, indicating that the encapsulation of CsoS2A was not mediated by interactions between the CsoS2-M in CsoS2B and CsoS2A. In addition, a higher degree of co-localization was found in cells expressing the CsoS2B-only shells than the CsoS2 shells (Figure 4c). This suggests that the latter has a reduced CsoS2A-loading capacity, which might be attributed to the competition between GFP-CsoS2A and endogenous CsoS2A for the limited binding sites on the shell inner surface. The CsoS2-C shells exhibited the lowest co-localization compared to the other three types of shell variants (Figure 4c), possibly due to the smallest size of the CsoS2-C shells, which likely results in a reduced number of docking sites for GFP-CsoS2A binding.

**Figure 4.**
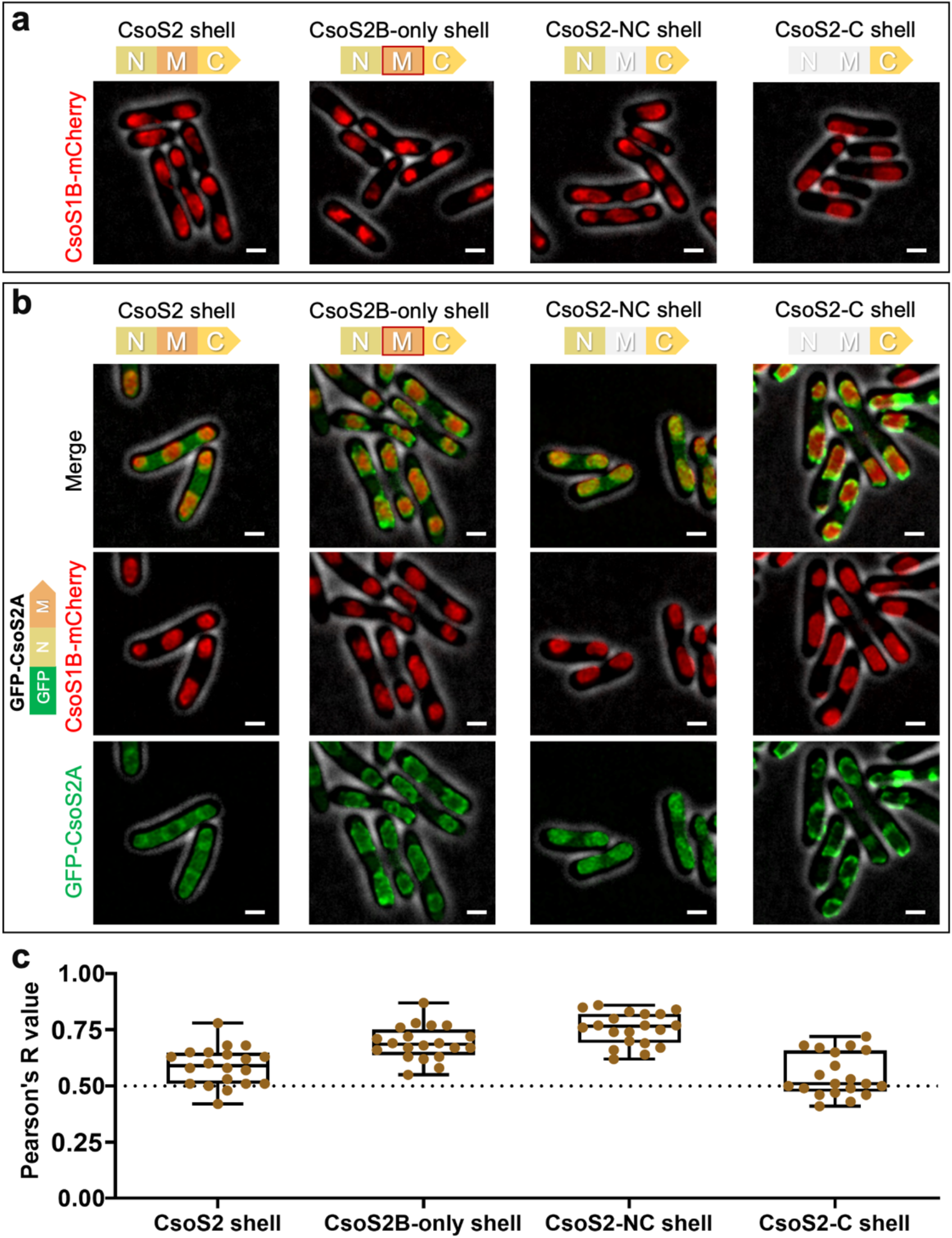
The encapsulation of CsoS2A is not mediated by interactions between CsoS2-M in CsoS2A and CsoS2B. **(a)** Confocal images of cells expressing CsoS2 shell, CsoS2B-only shell, CsoS2-NC shells, and CsoS2-C shells, respectively. CsoS1B was labeled by mCherry. Fluorescence foci (red) indicate shells. The CsoS2-M module with a red frame in the CsoS2B-only shell operon represents CsoS2-M without RFS. **(b)** Confocal images of cells co-expressing GFP-CsoS2A (green) with CsoS2 shell (red), GFP-CsoS2A with CsoS2B-only shell, GFP-CsoS2A with CsoS 2-NC shell, and GFP-CsoS2A with CsoS2-C shell. Scale bar: 1μm. **(c)** Colocalization analysis of GFP and mCherry fluorescence in **(b)** The Pearson’s R values for all the strains are: 0.59 ± 0.09 (CsoS2 shell); 0.69 ± 0.07 (CsoS2B-only shell); 0.76 ± 0.07 (CsoS2-NC shell); 0.55 ± 0.10 (CsoS2-C shell). Data are represented as mean ± SD. *n* = 20, representing the number of cells.

To further validate that the association between CsoS2-M of CsoS2A and the shell is responsible for CsoS2A encapsulation, we deleted the nucleotides encoding CsoS2-N from the plasmid expressing GFP-CsoS2A. This resulted in the generation of a GFP-fused CsoS2-M (GFP-S2M). We then co-expressed this construct with the mCherry-labeled shell variants. Confocal images revealed a comparable co-localization pattern of GFP and mCherry signals at the cell poles (Figure S6). This suggests that CsoS2-M within CsoS2A can mediate the encapsulation of CsoS2A into the shell, likely through the binding between the conserved [V/I/M][T/S]G motifs of CsoS2-M and shell hexamers (Julia et al., 2023; Ni et al., 2023; Zhou et al., 2024).

Additionally, the lack of the CsoS2A variant with one M-repeat in the purified CsoS2 (Mr=1) shells compared to the CsoS2 (Mr=3) shells suggests that the binding between a single M-repeat (M6) and the shell is insufficient to recruit CsoS2A(M6) into the shell (Figure S3b). To further examine the minimal number of M-repeats required for the encapsulation of CsoS2A, we generated a construct to produce a CsoS2 (Mr=2) shell variant that contains two M-repeats (M5-M6). SDS-PAGE showed the presence of CsoS2A(M5-M6) in the purified CsoS2 (Mr=2) shells (Figure S7), indicating that at least two M-repeats are required to ensure stable interactions between M-repeats and the shells for recruiting CsoS2A.

## Discussion

How the scaffolding protein CsoS2 governs the shell assembly and shell-cargo association is a key question not only in the fundamental studies of α-CB assembly but also in the bioengineering of CB-based nanostructures. This study provides a profound understanding of the role of CsoS2, particularly its middle region CsoS2-M, in controlling the size and curvature of α-CB shells in the absence of cargos. We show that CsoS2-M plays a dominant role in shaping the α-CB shell, possibly through strengthening the hexamer-hexamer association on both the facet-facet interfaces and flat shell facets distal from the shell vertices (Figure 5a). However, without CsoS2-C, neither CsoS2A nor CsoS2-M can independently orchestrate the assembly of the large α-CB shell (∼120 nm) (Figure 2b). More importantly, we show that the number of M-repeats in CsoS2-M plays a crucial role in determining the shell size and curvature, with a higher number of M-repeats resulting in enlarged polyhedral shells (Figure 5b).

**Figure 5.**
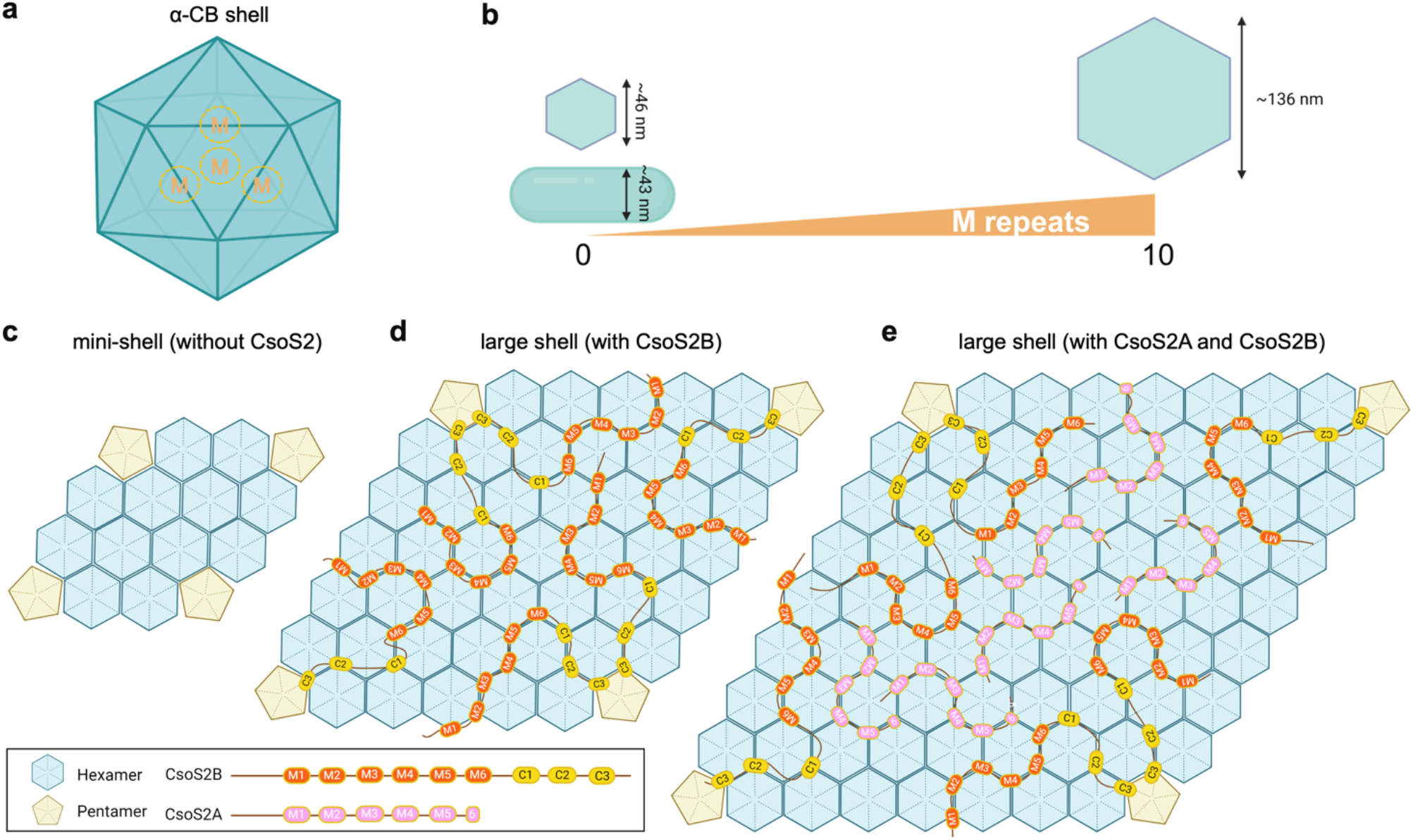
The role of CsoS2 in α-CB shell assembly. **(a)** The interaction sites of CsoS2-M on the α-CB shell. M represents CsoS2-M; the dashed circles indicate the anchoring sites of CsoS2-M on the shell. **(b)** The number of M-repeats defines the shell size and shape, with more M-repeats contributing to larger polyhedral shells. **(c)** Without CsoS2, shell proteins self-assemble into small facets, resulting in mini-shell formation. **(d)** With CsoS2B, shell proteins assemble into large facets, yielding large shells. The CsoS2-C connects the pentamer and the proximal hexamers; CsoS2-M reinforces hexamer-hexamer interfaces distal from the shell vertices. **(e)** With both CsoS2B and CsoS2A, shell proteins can form larger facets and larger shells, compared to CsoS2B-only shells. The CsoS2A adopts a flexible binding mode at the hexamer-hexamer interfaces distal from the shell vertices and consequently, contributes to larger shell formation.

Based on these observations, combined with our recent findings revealing the interactions between CsoS2-C with shell proteins and the cryo-EM structure of a small α-CB from *Prochlorococcus* (Ni et al., 2023; Zhou et al., 2024), we propose a model to elucidate the role of CsoS2 in shaping the α-CB shell. Without CsoS2, shell proteins could only self-assemble into mini-shells with a maximal size of ∼25 nm (Figure 5c). In the presence of CsoS2B, the CsoS2-C of CsoS2B attaches to the shell hexamers that surround the pentamer, while the flexible M-repeats of CsoS2-M extend outward, acting as a hinge to link neighbouring shell hexamers on both the flat facets away from shell vertices and the facet-facet interfaces. The increase in the content of CsoS2 M-repeats enables association with a higher number of shell hexamers. These structural features of the intrinsically disordered protein CsoS2 and the specific interactions of CsoS2-M with shell components facilitate the assembly of large shell facets and determine the tilt angles between adjacent shell facets, eventually promoting the formation of large polyhedral shells (Figure 5d).

The C-terminal truncated isoform of CsoS2, CsoS2A, has been proposed to solely coordinate Rubisco packaging within the α-CB, without attaching to the shell (Cai et al., 2015). Our results show that CsoS2A can be recruited into the empty shell and its encapsulation is driven by the interactions between the CsoS2-M of CsoS2A and the shell, instead of the interactions between CsoS2A and CsoS2B. Moreover, the integration of CsoS2A within the shell facilitates the assembly of larger shells, likely through stabilizing hexamer-hexamer association away from the curved shell vertices (Figure 5e). This ultimately leads to the formation of larger α-CB shells than the CsoS2B-only shells.

Our study offers experimental evidence that highlights the potential strategies for the natural design of CsoS2 in native α-CBs for defining and regulating the shell architecture to ensure the structural plasticity of α-CBs. The evolution of six M-repeats in native CsoS2 may be of physiological significance to enable the α-CB shell to form a large and stable structure, for the recruitment and packaging of a large number of Rubisco via CsoS2-N, while maintaining a stable polyhedral shape by the least six motifs in CsoS2-M (Figure 3c). Moreover, the multivalent interactions with shell proteins and the intrinsically disordered linker regions of CsoS2-M provide a means of modulating the assembly of shell proteins in a flexible mode and thereby modifying the α-CB shell structure (Figure 5d). This regulating capability is further strengthened by the presence of two isoforms of CsoS2 (Figure 5e). The structural plasticity of α-CBs mediated by CsoS2 may enable the host organisms to optimize their carbon fixation performance in response to varying environmental conditions.

In summary, this study provides mechanistic insights into the function of CsoS2, in particular, CsoS2-M, in shaping the formation and morphology of α-CB shells. Our findings advance the knowledge of the assembly principles of α-CBs and could inform rational design and reprogramming CBs and shell nanostructures for various biotechnological and biomedical applications.

## Methods

### Generation of constructs

Primers (Supplementary Table 1) for cloning genes and sequencing were ordered from Integrated DNA Technologies (US). All connections between genes and linearized vectors were achieved by Gibson assembly (Gibson assembly kit, New England BioLabs, UK). The shell-(*csoS2-NM*), shell-(*csoS2-NC*), shell-(*csoS2-C*), and shell-(Δ*csoS2*) operons were generated by deleting nucleotide sequences encoding CsoS2-C, CsoS2-M, CsoS2-C and CsoS2, respectively, from the synthetic shell operon derived from *Halothiobacillus neapolitanus* (Li et al., 2020). For the construction of variant shell operons expressing CsoS2 shells with varying numbers of M-repeats, the 6^th^ repeat (M6) containing RFS was constantly retained as the last M-repeat in the CsoS2-M to ensure the production of two isoforms. By contrast, for operons expressing CsoS2B shells, M6 was exclusively removed.

The shell-(*csoS2B-only*) operon was generated by replacing the *csoS2* gene, in the shell operon, with the *csoS2B* gene (Chaijarasphong et al., 2016), respectively. To construct fluorescence-tagged shell operons, the gene encoding mCherry was fused to the C-terminus of the *csoS1B* gene in various shell operons. The enhanced *gfp* gene, with the nucleotides encoding CsoS2-M, or CsoS2A fused at the C-terminus, was cloned into pCDFDueT-1 linearized by NcoI and XhoI. These constructs were placed under the control of a pTrc promoter, resulting in the generation of the pCDF-GFP-CsoS2M and pCDF-GFP-CsoS2A vectors, respectively. All these constructs were verified by PCR and DNA sequencing and transformed into *E. coli* Top10 and BW25113 cells.

### Expression and Isolation of α-CB shells

*E. coli* BW25113 strains containing various *cso* vectors were cultivated at 37 °C in Lysogeny Broth (LB) medium containing 100 μg mL^−1^ ampicillin. The expression of these vectors was induced by L-Arabinose (1 mM, final concentration) once the cells reached an early log phase (OD_600_ = 0.6). Cells were grown at 25 °C for 16 hours with constant shaking and then were harvested by centrifugation at 5000g for 6 minutes. The cell pellets were washed with TEMB buffer (10 mM Tris-HCl, pH = 8.0, 1 mM EDTA, 10 mM MgCl_2_, 20 mM NaHCO_3_) and resuspended in TEMB buffer supplemented with 10% (v/v) CelLytic B cell lysis reagent (Sigma-Aldrich) and 1% protein inhibitor cocktail (100×) (Sigma-Aldrich).

The CsoS2 shells and CsoS2B-only shells were purified following the standard shell purification procedures at 4 °C. The cell suspensions were lysed by sonication, and cell debris was removed by centrifugation at 12,000 *g* for 10 minutes, followed by centrifugation at 50,000 *g* for 30 minutes to enrich shells. The pellets were resuspended in TEMB buffer and then loaded onto sucrose gradients (10−50%, w/v) followed by ultracentrifugation (Beckman, XL100K ultracentrifuge) at 105,000 *g* for 30 minutes. The CsoS2-C shells and Δ*csoS2* shells were purified following the mini-shell purification protocol described previously (Ni et al., 2023).

To isolate shells with varying numbers of M-repeats, we slightly modified the purification procedures from the standard protocol. These adjustments were made to ensure that the shells were distributed within the 10−50% sucrose fractions. For shells with no more than five M-repeats, the purification process involved removing the cell debris, then subjecting the supernatants to centrifugation at 50,000 *g* for 30 minutes, followed by ultracentrifugation at 105,000 *g* for 1 hour. Shells with seven M-repeats were purified using the standard purification protocol. For shells with ten M-repeats, after removing the cell debris, the supernatants were subjected to centrifugation at 50,000 *g* for 30 minutes, followed by ultracentrifugation at 105,000 *g* for 15 minutes. Each sucrose fractions were collected and stored at 4 °C.

### Expression and Isolation of GFP-loaded α-CB shells

*E. coli* BW25113 strains co-expressing shells and GFP-tagged proteins were cultivated at 37 °C in lysogeny broth (LB) medium containing 100 μg mL^−1^ ampicillin and 50 μg mL^−1^ spectinomycin. The expression of GFP-fused cargos was induced by the addition of 0.1 mM IPTG at OD_600_ = 0.6. After 4 hours of induction of the expression of GFP fusions, the shell expression was induced by 1 mM L-arabinose, and cells were then grown at 25 °C for 16 hours. The isolation of GFP-incorporated shells was purified following the standard purification protocol described above.

### SDS-PAGE analysis

SDS−PAGE was performed following the procedure described previously. Briefly, 40 μg of total protein was loaded into each well of 16% polyacrylamide gels and stained with Coomassie Brilliant Blue R250.

### Transmission electron microscopy

Thin-section transmission electron microscopy (EM) was performed to visualize the reconstituted shell structures in *E. coli* strains (Fang et al., 2018). Isolated shell structures were characterized using negative staining EM. Images were recorded using an FEI Tecnai G2 Spirit BioTWIN transmission electron microscope equipped with a Gatan Rio 16 camera. Image analysis was carried out by using ImageJ software. The shell diameter data was randomly collected from 100 shell particles on EM images. The diameter of each polyhedral shell particle was measured by drawing diagonals three times from various angles, all intersecting at the same center point, using ImageJ software, and the resulting measurements were then averaged. The irregularity degree was determined by calculating the ratio of the standard deviation to the average of three diagonal measurements for each shell.

### Confocal microscopy

Overnight-induced *E. coli* cells were immobilized by drying a droplet of cell suspension onto LB agar pads as described previously (Yang et al., 2022). Blocks of agar with the cells absorbed onto the surface were covered with a cover slip and placed under the microscope. Laser-scanning confocal fluorescence microscopy imaging was performed on a Zeiss Elyra 7 with Lattice SIM^2^ microscope equipped with a 63×/1.4 NA oil immersion objective, excitation wavelength at 488 and/or 561 nm. GFP and mCherry fluorescence were detected at 500– 520 nm and 660–700 nm, respectively. Live-cell images were recorded from at least three different cultures. All images were captured with all pixels being below saturation. Image analysis was carried out using ImageJ software.

### Statistics and reproducibility

All experiments reported here were performed at least three times independently and at least three biological repeats were performed for each experiment.

