## Supplementary Information for "Uncovering the roles of the scaffolding protein CsoS2 in mediating the assembly and shape of the α-carboxysome shell"

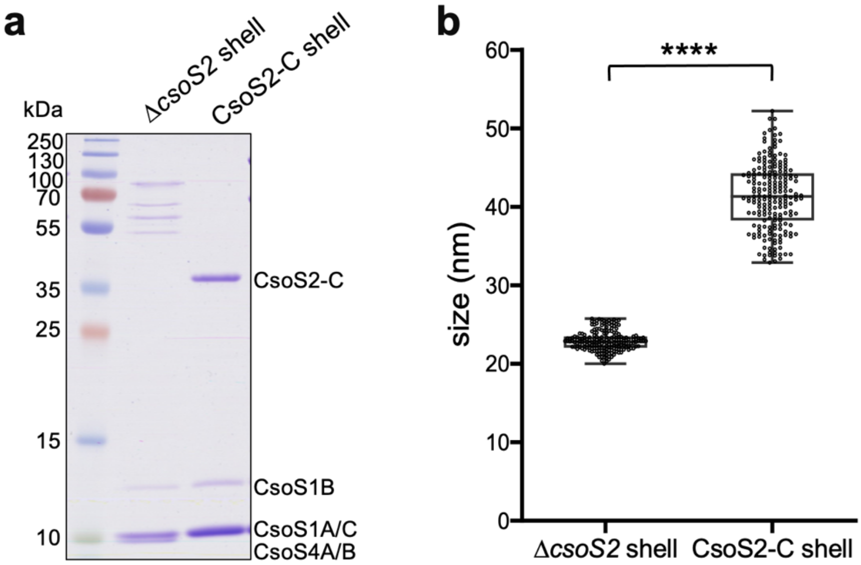


**Figure S1. Characterization of Δ*csoS2* shells and CsoS2-C shells. (a)** SDS-PAGE of purified Δ*csoS2* shells and CsoS2-C shells. **(b)** Size comparison between Δ*csoS2* shells and CsoS2-C shells. **** $p < 0.0001$ ($n = 100$, two-tailed unpaired t-test). Box plots indicate the median (middle line in the box), 25th percentile (bottom line of the box), and 75th percentile (top line of the box).

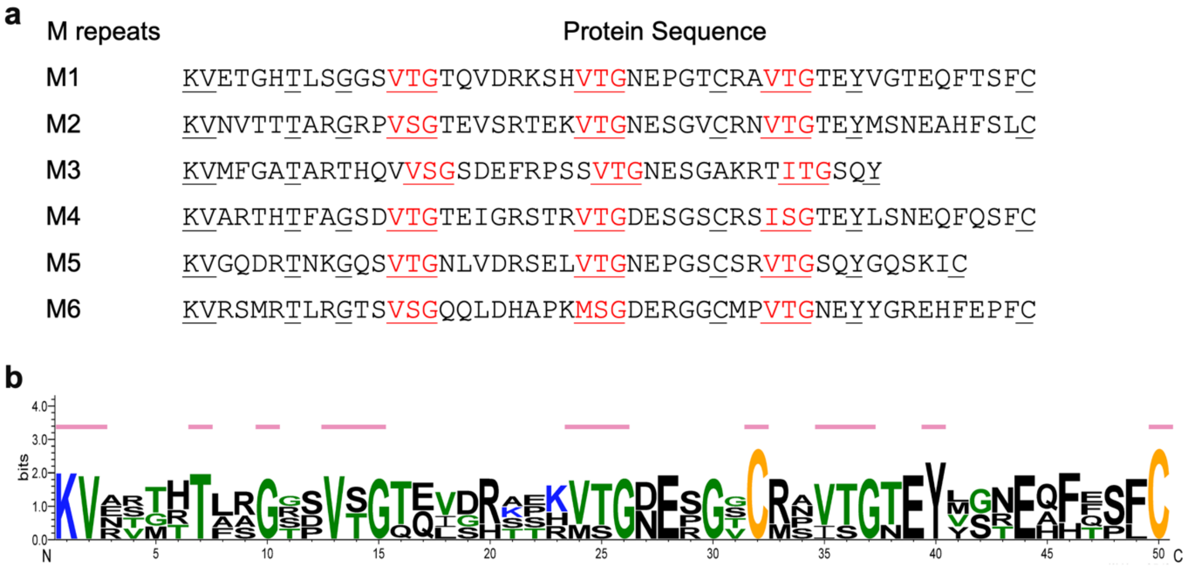


**Figure S2. Conservation analysis of M-repeats. (a)** Protein sequences of each middle region repeat. Conservative residues are underlined. **(b)** Sequence logo for M-repeats presented using Weblogo 3. Key conserved residues are indicated by pink lines.

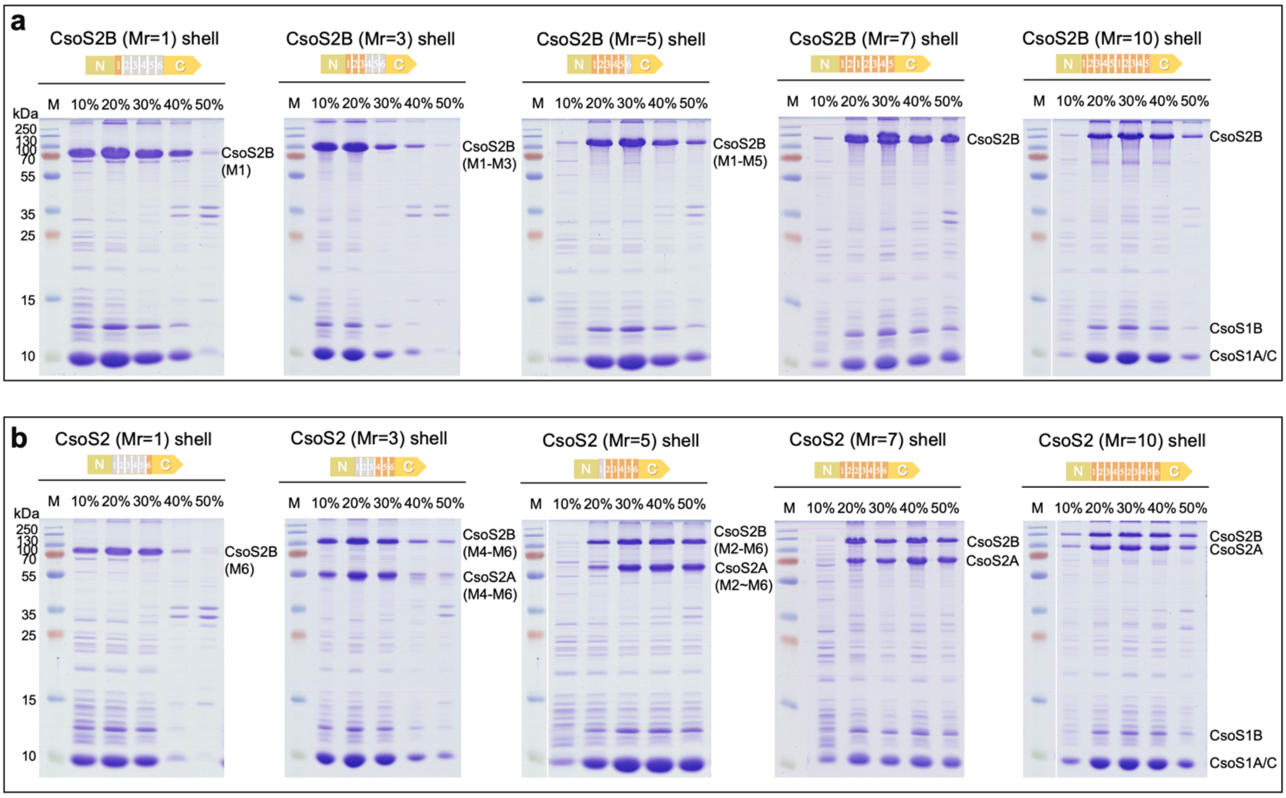


**Figure S3. SDS-PAGE of purified CsoS2B and CsoS2 shells with variable M-repeats (the deleted regions are colored grey). (a)** SDS-PAGE of CsoS2B shells with variable M-repeats (Mr) in 10-50% sucrose fractions. **(b)** SDS-PAGE of CsoS2 shells with variable M-repeats in 10-50% sucrose fractions.

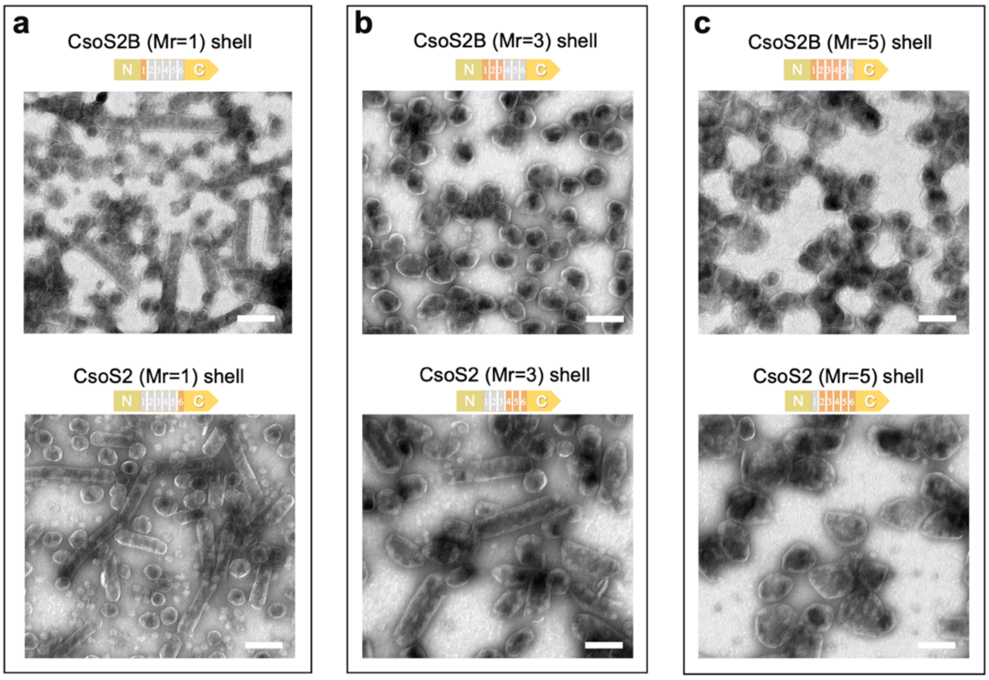


**Figure S4. Shell curvature correlates with the increase in the number of M-repeats (the deleted regions are colored grey). (a)** EM of purified CsoS2B (Mr=1) shells (top) and CsoS2 (Mr=1) shells (bottom). **(b)** Purified CsoS2B (Mr=3) shells (top) and CsoS2 (Mr=3) shells (bottom). **(c)** Purified CsoS2B (Mr=5) shells (top) and CsoS2 (Mr=5) shells (bottom). Samples were taken from the 40% sucrose fractions. Scale bar, 100nm.

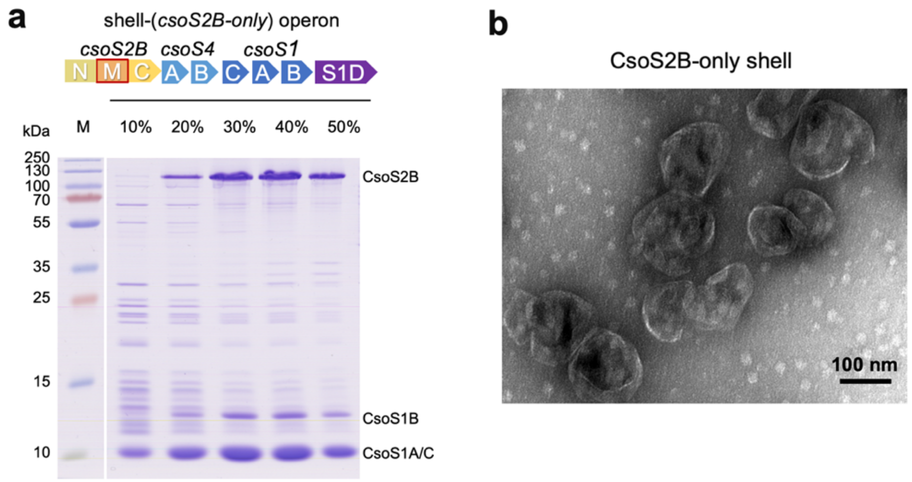


**Figure S5. Characterization of CsoS2B-only *shells*. (a)** SDS-PAGE of proteins purified from cells expressing shell-(*csoS2B-only*) operon in the 10-50% sucrose fractions. **(b)** EM of purified CsoS2B-only shells in the 30% sucrose fraction.

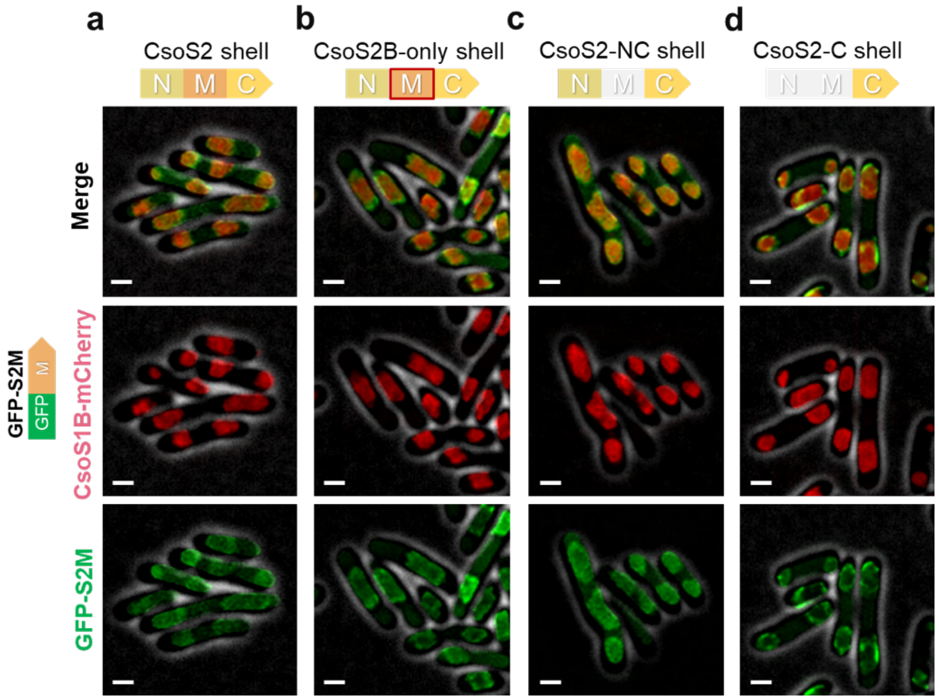


**Figure S6. The encapsulation of CsoS2A is mediated by the interaction between the middle region of CsoS2A with shell proteins. (a-d)** Confocal images of cells co-expressing GFP-S2M (green) with CsoS2 shells (red) (a); GFP-S2M with CsoS2B-only shells (b); GFP-S2M with CsoS2-NC shells (c); GFP-S2M with CsoS2-C shells (d)**.** Scale bar: 1 µm.

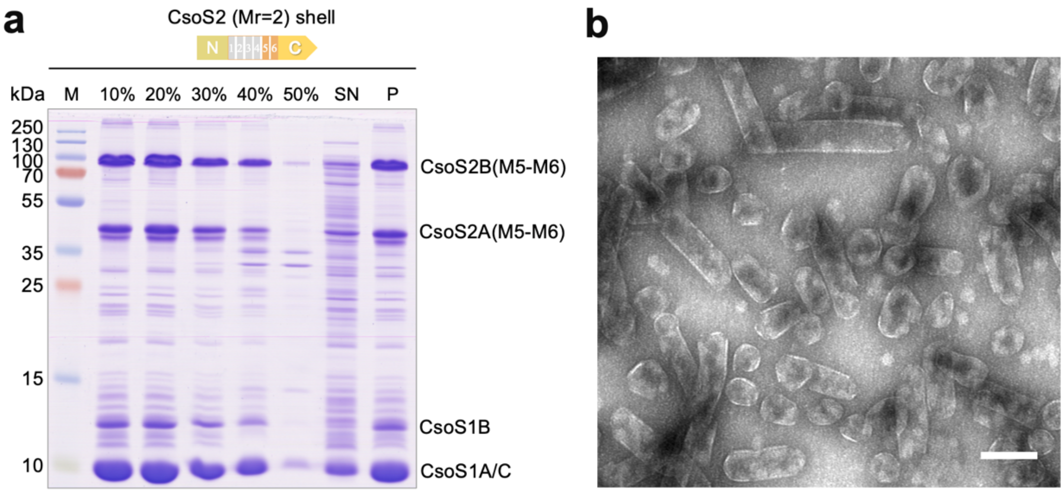


**Figure S7. SDS-PAGE and EM analysis of purified CsoS2 (Mr=2) shells. (a)** SDS-PAGE of CsoS2 shells with two M-repeats in 10-50% sucrose fractions. **(b)** EM of purified CsoS2 (Mr=2) shells in 20% sucrose fraction. Scale bar, 100 nm.

**Supplementary Table 1. Primers used in this study.** The overlapping sequences for Gibson assembly are underlined.

| Primer | Sequence (5'-3') | Description |
|---|---|---|
| pBAD-EcoRI-FW | cgaagcttacgtagaacaaaaactc | Construction of shell-($\Delta$*csoS2*) operon |
| pBAD-NcoI-RV | ggttaattcctcctgttagc | |
| pBAD-RBS-CsoS4A-FW | ggctaacaggaggaattaactttgagcgttcaggcgcag | |
| pBAD-CsoS1D-RV | tgttctacgtaagcttcgttattagaacccttcagcgcgac | |
| NcoI-pBAD-S2-C-FW | ggctaacaggaggaattaaccatgccgttttgtacgagcaccccagag | shell-(*csoS2-C*) operon |
| EcoRI-pBAD-S1D-RV | ttgttctacgtaagcttcg | |
| CsoS2(NC)-FW | gggcaacgcagcgccaaaaacgagcaccccagagcccgaa | shell-(*csoS2-NC*) operon |
| CsoS2(N)-RV | ttttggcgctgcgttgccc | |
| CsoS2(NM)-FW | ttaagtaaagtgtaacgatctttgagcg | shell-(*csoS2-NM*) operon |
| CsoS2(NM)-RV | cgctcaaagatcgttacactttacttaatcaacaaaacggttcgaaatgttcacg | |
| pBAD-NcoI-S2-FW | ggctaacaggaggaattaaccatgccttcacagtcaggaatg | shell-(*csoS2-only*) operon |
| CsoS4A-CsoS2-RV | cgtagtactcattaccggtaactggcatacatccacctcgttcgtcaccggacatc | |
| A147 C1-FW | acgagcaccccagagcccgaa | *csoS2B(Mr=1)* shell operon |
| A146 ko(M2-M6)-RV | ttcgggctctggggtgctcgtcgtcgcatttggcttggg | |
| A149 ko (M1-M5)-FW | gggcaacgcagcgccaaaaggtggtggcgtgggaaaagtgc | *csoS2(Mr=1)* shell operon |
| A75 S2(N)-RV | ttttggcgctgcgttgccc | |
| A218 KO(M1-M4)-FW | gggcaacgcagcgccaaaagacacaaaacctcaacgcagccc | *csoS2(Mr=2)* shell operon |
| A186 pBAD-RV | ggttaattcctcctgttagc | |
| A158 M3-RV | ttcgggctctggggtgctcgttgcaggtgctccgttgatcgtgagt | *csoS2B(Mr=3)* shell operon |
| A162 ko M3-FW | gggcaacgcagcgccaaaagcagacgaaggtcttgcgcgact | *csoS2(Mr=3)* shell operon |
| A148 ko M6-RV | ttcgggctctggggtgctcgttcccacgccaccaccgcagatttt | *csoS2B(Mr=5)* shell operon |
| A150 ko M1-FW | gggcaacgcagcgccaaaaaccagccccaagccaaatgcg | *csoS2(Mr=5)* shell operon |
| A195 M1-FW | gggcaacgcagcgccaaaaaaggttgaaaccggtcacaccct | *csoS2B(Mr=7)* shell operon |
| A196 (M1)-M2-RV | gtgtgaccggtttcaaccttatccgcttgtgaaggctttgc | |
| A197 (M2)-M2-RV | ccgttgtggtcacattgaccttcgtatccgcttgtgaaggctttgc | *csoS2(Mr=7)* shell operon |
| A159 N4-M1-FW | gggcaacgcagcgccaaaaaaggttgaaaccggtcacaccct | *csoS2B(Mr=10)* shell operon |
| A160 M1-FW | aaggttgaaaccggtcacaccct | |
| A161 M5-RV | agggtgtgaccggtttcaacctttcccacgccaccaccgcagatttt | |
| A164 M2-FW | acgaaggtcaatgtgaccacaacgg | *csoS2(Mr=10)* shell operon |
| A163 M2-M5-RV | gtggtcacattgaccttcgttcccacgccaccaccgcagatttt | |
| A202 GS-mCherry-FW | ggtggtagcggtggtagtagcaagggcgaggaggat | Fuse the *mCherry* gene to the C-terminus of *csoS1B* |
| A203 S1D-mCherry-RV | gcgcatcttccctactagacattacttgtacagctcgtccatgc | |
| A204 GS-S1B-RV | actaccaccgctaccaccgctattcagatttgcgatacacc | |
| A190 S57 pCDF(NcoI)-pTrc-FW | tgtttaactttaataaggagatatacgtaaatcactgcataattcgtgt | pCDF-GFP-S2M and pCDF-GFP-CsoS2A |
| A191 GS-GFP-RV | actaccaccgctaccaccctt | |
| A192 GFP-S2(M-FW) | aagggtggtagcggtggtagtggcacagcaccttcctgcaag | |
| A193 pACYC-S2A-RV | gcagcggtttctttaccagactcagcgtacgtcctttggg | |
| A194 GFP-S2A-FW | aagggtggtagcggtggtagtccttcacagtcaggaatgaat | |
